# Sustained fitness gains and variability in fitness trajectories in the long-term evolution experiment with *Escherichia coli*

**DOI:** 10.1101/027391

**Authors:** Richard E. Lenski, Michael J. Wiser, Noah Ribeck, Zachary D. Blount, Joshua R. Nahum, J. Jeffrey Morris, Luis Zaman, Caroline B. Turner, Brian D. Wade, Rohan Maddamsetti, Alita R. Burmeister, Elizabeth J. Baird, Jay Bundy, Nkrumah A. Grant, Kyle J. Card, Maia Rowles, Kiyana Weatherspoon, Spiridon E. Papoulis, Rachel Sullivan, Colleen Clark, Joseph S. Mulka, Neerja Hajela

## Abstract

Many populations live in environments subject to frequent biotic and abiotic changes. Nonetheless, it is interesting to ask whether an evolving population’s mean fitness can increase indefinitely, and potentially without any limit, even in a constant environment. A recent study showed that fitness trajectories of *Escherichia coli* populations over 50,000 generations were better described by a power-law model than by a hyperbolic model. According to the power-law model, the rate of fitness gain declines over time but fitness has no upper limit, whereas the hyperbolic model implies a hard limit. Here, we examine whether the previously estimated power-law model predicts the fitness trajectory for an additional 10,000 generations. To that end, we conducted more than 1100 new competitive fitness assays. Consistent with the previous study, the power-law model fits the new data better than the hyperbolic model. We also analysed the variability in fitness among populations, finding subtle, but significant, heterogeneity in mean fitness. Some, but not all, of this variation reflects differences in mutation rate that evolved over time. Taken together, our results imply that both adaptation and divergence can continue indefinitely— or at least for a long time—even in a constant environment.

## 1. Introduction

In nature, the process of adaptation by natural selection appears inexhaustible and openended in its creativity [1]. Such sustained adaptation is usually thought to occur in response to changing environmental conditions, including those produced by the evolution of other organisms with which the focal population interacts. As a corollary, it is also often presumed that evolving organisms must eventually “run out” of ways to improve, absent changes in their environment. In the parlance of adaptive landscapes, a population will, sooner or later, reach a local fitness peak [2,3].

Wiser et al. [4] challenged the presumption that there must be an upper bound to organismal fitness. They measured the fitness trajectories over 50,000 generations for *Escherichia coli* populations in the long-term evolution experiment (LTEE). They compared the fit of two simple models—a hyperbolic model and a power-law model—that both predict decelerating rates of fitness improvement, but only the former has an upper limit, or asymptote. The power-law model, by contrast, predicts that the logarithm of fitness will increase with the logarithm of time, a relationship that has no asymptote. Both models fit the observed fitness trajectories well, but the power-law model fit much better. Moreover, if truncated datasets (e.g., from only the first 20,000 generations) were used to predict the subsequent trajectories, the hyperbolic model consistently underestimated the extent of future improvement, whereas the power-law model accurately predicted the changes seen in later generations. Despite having no upper limit, but owing to its logarithmic dependence on time, the power law did not lead to absurd predictions that would seem to violate physical constraints. Indeed, when the power-law model was extrapolated millions of generations into the future, the predicted fitness levels correspond to growth rates that are within the range that some bacterial species can achieve under optimal conditions.

In addition, Wiser et al. [4] presented a dynamical model of fitness evolution for large asexual populations that also generated a power-law relationship. The dynamical model included two particularly important phenomena that have been documented in the LTEE—clonal interference, in which contemporaneous lineages with different beneficial mutations compete for fixation [5,6]; and diminishing-returns epistasis, in which the fitness advantage of beneficial mutations tends to be smaller on more-fit genetic backgrounds [7]. That model also predicted that several populations that evolved hypermutable phenotypes [8] early in the LTEE would show faster rates of fitness improvement, and this prediction was confirmed.

Here, we perform competition assays using population samples from 40,000, 50,000, and 60,000 generations to test whether the power-law model’s predictions continue to hold. Because the magnitude of fitness changes are predicted to become increasingly small—and the curvature of the fitness trajectory increasingly subtle—as time goes by, we performed more than 1100 competitions to assess the fit of the model. This extensive replication also allowed us to estimate with some precision the among-population variance component for mean fitness (i.e., the variation above the level attributable to the measurement error in replicate assays). From previous studies, we know that one of the 12 LTEE populations followed a different path from the others, when it evolved the ability to grow on citrate in the medium (included as a chelating agent), which neither the ancestor nor any of the other populations can exploit [9,10]. We also know that the populations that became hypermutable achieved a boost in their fitness trajectories [4]. Early in the LTEE, before these exceptional cases had evolved, previous estimates of the among-population variance indicated subtle but significant heterogeneity in mean fitness [11]. However, it was not known whether that variation was transient or would be sustained over the long term. Our results add support to the hypothesis that adaptive evolution is unbounded even in a constant environment and show that the subtle divergence of the populations’ fitness trajectories continues unabated as well.

## 2. Materials and Methods

### (a) Strains

The *E. coli* long-term evolution experiment (LTEE) is described in detail elsewhere [11,12]. Our analyses used whole-population samples from nine of the 12 LTEE populations taken at three time points: 40,000, 50,000, and 60,000 generations (Table S1), for a total of 27 samples. We excluded three populations (Ara−2, Ara−3, and Ara+6) that no longer make colonies that can be reliably counted in the standard fitness assays or that evolved the ability to consume the citrate in the culture medium, which also precludes using the standard assays [4]. Of the nine populations used in our study, three evolved hypermutable phenotypes: Ara−1, Ara−4, and Ara+3 [8,13]. In addition to the whole-population samples, we used two clones as common competitors, REL10948 and REL11638 [4]. The former is an Ara^−^ clone isolated from population Ara−5 at 40,000 generations; the latter is a spontaneous Ara^+^ mutant of that clone (Table S1). The Ara marker serves to distinguish competitors during fitness assays, as Ara^−^ and Ara^+^ cells make red and white colonies, respectively, on tetrazolium-arabinose (TA) indicator plates. This marker has been shown to be selectively neutral under the glucose-limited conditions of the LTEE [14,15]. The use of an evolved strain as a common competitor should improve the precision of estimates when the fitness differential between the evolved and ancestral types is large [16]. We also used the LTEE’s ancestral strain, REL606 [11,17], and three clones from populations Ara+4 and Ara+5 at 60,000 generations (Table S1) in assays to estimate mutation rates.

### (b) Culture conditions

The culture conditions for the LTEE are described elsewhere [11,12]. In brief, each population is maintained by transferring 0.1 mL of culture into 9.9 mL of fresh medium every 24 h. The medium, called DM25, contains 25 μg/mL glucose, which is the limiting resource. Cultures are kept at 37°C in a shaking incubator for aeration. The 100-fold dilution and regrowth allow ~6.6 generations per day. Every 500 generations (75 days), samples of each population are stored in 10% glycerol at −80°C; the bacteria in those samples remain viable and are available for later study.

### (c) Fitness assays

The fitness assays followed the same procedures as used in the first experiment reported by Wiser et al. [4]. In brief, fitness was measured in the same environment as used in the LTEE by competing a population sample against a reference strain, either REL10948 or REL11638, with the opposite Ara marker state. Prior to the start of the assay, competitors were removed from the freezer and separately acclimated to the culture medium and other conditions described above. The competitors were then mixed at an equal volumetric ratio, and a sample was spread on a TA plate to estimate their initial abundances based on colony counts. The mixed competition cultures were propagated for three days by 1:100 daily dilutions into fresh medium and, at the end of the third day, another sample was spread on a TA plate to estimate the final abundances of the two competitors. From the initial and final counts of each type, and taking into account the dilution factors, we calculated for each competitor its realized growth rate during the assay. We then calculated fitness as the ratio of the evolved population’s growth rate to that of the reference strain. The assays were performed in 42 complete blocks of 27 assays each. Twenty authors performed one block each, and one author (M.J.W.) performed 22 blocks.

### (d) Missing values and outliers

Eight of the 1,134 assays failed to yield fitness estimates owing to procedural errors. In addition to the missing values, we screened the remaining estimates for outliers as follows. We first log-transformed the estimates to make random deviations symmetric around the mean. We then computed the mean and standard deviation of the transformed estimates for each of the 27 population samples, and we converted each transformed estimate to a *z* score by taking the absolute value of its deviation from the mean and dividing by the standard deviation. Assuming normality, one expects ~0.3% of the *z* scores to be greater than 3 and only ~0.01% to be greater than 4. Thus, one would expect to see among the 1,126 estimates only a few *z* scores greater than 3 and none or at most one above 4. However, 13 *z* scores were over 3, including 5 that exceeded 4. There were also 44 scores that fell between 2 and 3; assuming normality, we expect 48 such values, and so there are no excess deviations in this range. Based on this analysis, we eliminated the estimates that produced the 13 most extreme *z* scores from our analyses, leaving a total of 1,113 values. The number of estimates ranged from 39 to 42 for the 27 population samples (Table S2). We also computed quantities that describe changes in fitness over time, which require estimates from multiple time points. Owing to the missing values and outliers, and because we calculated these changes using fitness estimates from the same population and block, we often had fewer estimates for these quantities. In the Results, we describe the criteria used for including data in each analysis.

### (e) Fluctuation tests

We performed fluctuation tests [18] to compare the mutation rates of the ancestor and three 60,000-generation clones from each of two populations, Ara+4 and Ara+5. The bacteria were revived by inoculating 15 μL of frozen stock into 10 mL of LB broth, then incubating the cultures overnight at 37°C in a shaking incubator for aeration. Each resulting stationary-phase culture was then diluted 10,000-fold into 9.9 mL of DM25 and incubated for 24 h at 37°C with aeration. From each DM25 culture, we transferred 150–600 cells to 24 replicate 600-μL cultures of DM250 (containing 250 μg/mL glucose) in 96-well plates, which were incubated for 24 h at 37°C. Samples from six replicate cultures were then diluted and spread on TA plates to estimate the average number of cells, and 200-μL aliquots (one-third of the total volume) from all 24 cultures were spread onto LB plates supplemented with either 100 μg/mL rifampicin or 30 μg/mL nalidixic acid. The antibiotic-containing plates were incubated at 37°C for 48 h, and the number of plates with one or more resistant colonies was counted. The mutation rate, μ, was estimated using the *p*_0_ method [18] as −ln(*p*0)/*N*, where *p*_0_ is the proportion of replicate cultures without any mutants and *N* is the number of cells tested per replicate culture. If none of the replicates in a test produced any resistant mutant, then the mutation rate was estimated, by convention, using 0.5 as the number of positive replicates.

### (f) Statistical analyses

Population and block are random factors in ANOVAs. Statistical analyses were performed using R version 3.0.2 [19].

## 3. Results

### (a) Fitness continues to increase

In all nine populations tested, the estimated mean fitness increased over both the 40,000- to 50,000- and the 50,000- to 60,000-generation intervals (Figure 1, Table S2). The probability that all nine populations would, by chance, yield monotonically increasingly point estimates across three time points is (1/3 x 1/2)^9^ < 10^−7^.

**Figure 1.**
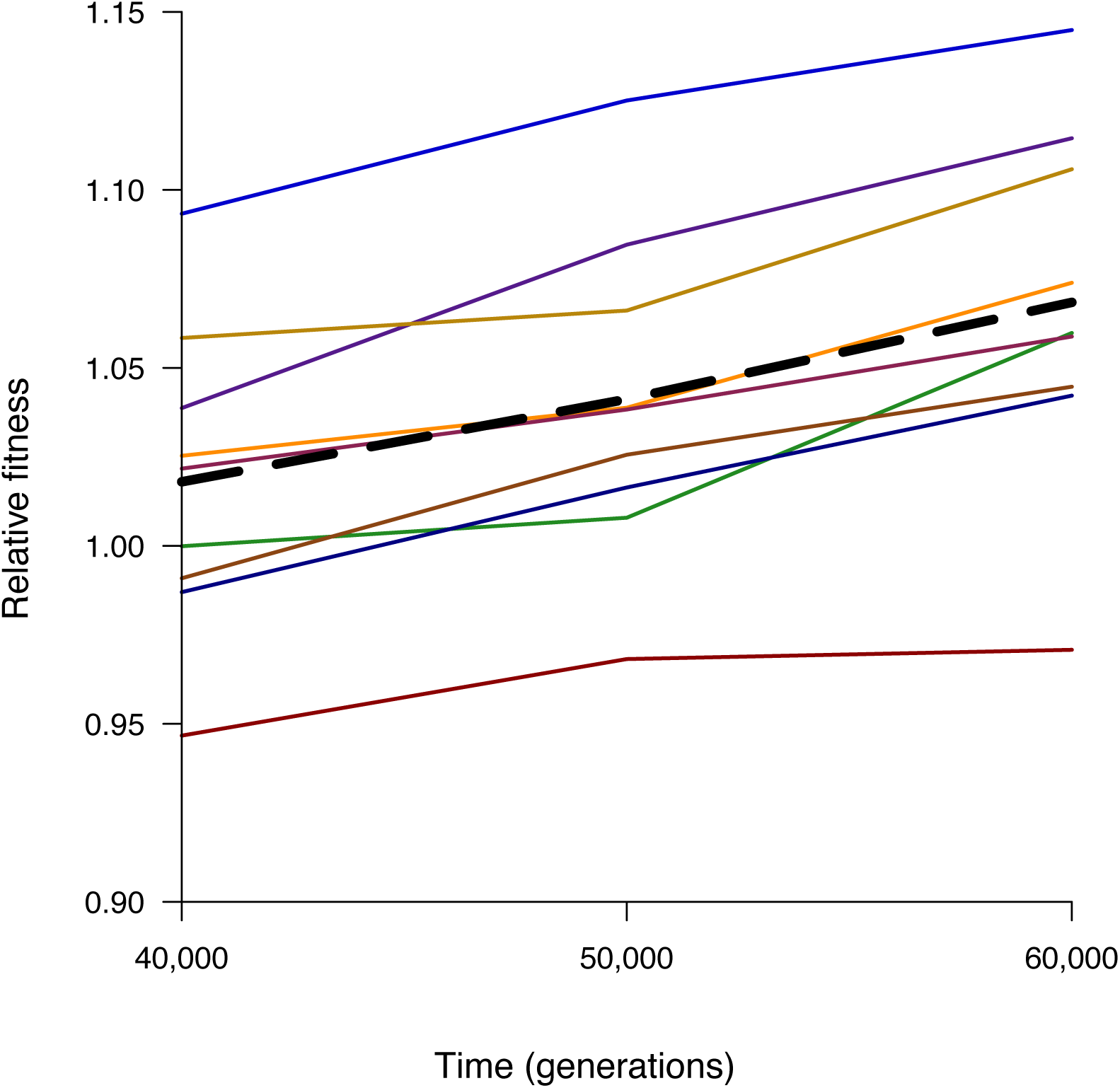
Trajectories of mean fitness for nine *E. coli* populations from the LTEE (thin lines) and grand mean fitness (thick dashed line). Data from Table S2. (Online version in colour.)

When each population is analysed individually (Table 1), the cumulative increase between 40,000 and 60,000 generations is highly significant (all *p* < 0.001). Six populations showed significant fitness gains over both component intervals, while three had significant gains in only one interval (all *p* < 0.05).

**Table 1.** Changes in mean fitness between 40,000 and 60,000 generations in the LTEE populations.

| **Generations** |  | **40,000 to 60,000** |  |  | **40,000 to 50,000** |  |  | **50,000 to 60,000** |  |  |
| --- | --- | --- | --- | --- | --- | --- | --- | --- | --- | --- |
| **Population** | **N** | **Mean** | **SD** | ***p*** | **Mean** | **SD** | ***p*** | **Mean** | **SD** | ***P*** |
| **Ara+1** | 39 | 0.0250 | 0.0383 | 0.0001 | 0.0227 | 0.0410 | 0.0007 | 0.0023 | 0.0359 | <u>0.3436</u> |
| **Ara+2** | 38 | 0.0527 | 0.0416 | <0.0001 | 0.0192 | 0.0533 | <u>0.0164</u> | 0.0335 | 0.0686 | 0.0023 |
| **Ara+3** | 38 | 0.0527 | 0.0489 | <0.0001 | 0.0335 | 0.0391 | <0.0001 | 0.0192 | 0.0502 | <u>0.0120</u> |
| **Ara+4** | 41 | 0.0603 | 0.0437 | <0.0001 | 0.0085 | 0.0342 | <u>0.0598</u> | 0.0519 | 0.0468 | <0.0001 |
| **Ara+5** | 40 | 0.0551 | 0.0410 | <0.0001 | 0.0297 | 0.0329 | <0.0001 | 0.0254 | 0.0420 | 0.0002 |
| **Ara-1** | 41 | 0.0756 | 0.0641 | <0.0001 | 0.0464 | 0.0528 | <0.0001 | 0.0292 | 0.0771 | <u>0.0099</u> |
| **Ara-4** | 39 | 0.0472 | 0.0425 | <0.0001 | 0.0059 | 0.0490 | <u>0.2289</u> | 0.0414 | 0.0483 | <0.0001 |
| **Ara-5** | 40 | 0.0541 | 0.0329 | <0.0001 | 0.0348 | 0.0272 | <0.0001 | 0.0193 | 0.0332 | 0.0003 |
| **Ara-6** | 42 | 0.0371 | 0.0267 | <0.0001 | 0.0165 | 0.0311 | 0.0007 | 0.0205 | 0.0329 | 0.0001 |

Based on N blocks for which fitness values were obtained at all three time points. All *p*-values were calculated using one-tailed *t*-tests, given the expectation that fitness should increase. All tests are significant ( $p < 0.05$ ) except three that were non-significant based on individual tests (double underline) and three others that were non-significant after a table-wide sequential Bonferroni correction (single underline).

One can apply a table-wise sequential Bonferroni correction [20,21] to the data in Table 1 to account for the facts that (i) we performed 27 significance tests in total and (ii) the sets of 3 tests for each population are not independent. However, the results hardly change—only 3 of the 24 tests that were individually significant at *p* < 0.05 become non-significant using this more conservative approach.

### (b) Gains more consistent with power-law model than with hyperbolic model

Wiser et al. [4] presented several lines of evidence that fitness trajectories of the LTEE populations are better described by a power-law model than by a hyperbolic model. Both models were fit to trajectories based on competitions between evolved population samples and their genetically marked ancestors using data through 50,000 generations. The best fit of the hyperbolic model gave *w* = 1 + 0.7007 *t* / (t + 4431), where *w* is the grand mean fitness relative to the ancestor and *t* is time in generations. This equation predicts grand mean fitness values relative to the ancestor of 1.6308, 1.6437, and 1.6525 at 40,000, 50,000, and 60,000 generations, respectively. For the power-law model, the best fit yielded *w* = (0.00515 *t* + 1)^0.0950^, which predicts values of 1.6597, 1.6951, and 1.7246 at the same time points.

In our study, the evolved population samples competed against a high-fitness strain (a 40,000-generation clone from one of the LTEE populations), not against their much less fit ancestor. This high-fitness strain was used so that the fitness differences between the competitors were much smaller, which allowed the competitions to run for three days and, in turn, provided more precise estimates of small differences in relative fitness. Therefore, to compare the fitness gains estimated in our study with the predictions of the models from Wiser et al. [4], we needed to analyse the *changes* in fitness, rather than fitness itself. The hyperbolic model predicts a proportional fitness increase of 0.79% (i.e., 1.6437/1.6308 - 1) from 40,000 to 50,000 generations, and it predicts an increase of 0.54% from 50,000 to 60,000 generations; the predicted increase over the two intervals combined is 1.33%. The power-law model predicts increases of 2.13% and 1.74% over the same two intervals, with a cumulative gain of 3.91%.

We can test the actual fitness gains against these two sets of predictions using the means for the nine populations (Table 2), where the independently evolving populations serve as the appropriate unit of replication in the comparisons. The observed fitness gains significantly exceed the gains predicted by the hyperbolic model over all three intervals (all *p* < 0.01). This outcome agrees with the pattern reported by Wiser et al. [4] when they truncated their 50,000-generation dataset and used shorter-duration subsets to predict the future trajectory of mean fitness—namely, the hyperbolic model consistently underestimated the potential for future gains. The observed gains are much closer to those predicted by the power-law model, although the difference between the model and observations was marginally significant (*p* = 0.035) over the entire 40,000- to 60,000-generation interval (Table 2). The observed improvement during that time was 5.1%, whereas the power-law model predicted a gain of 3.9%.

**Table 2.** Comparison between the observed changes in the grand-mean fitness and those predicted by the hyperbolic and power-law models.

| **Generations** | **40,000 to 60,000** | **40,000 to 50,000** | **50,000 to 60,000** |
| --- | --- | --- | --- |
| **Grand Mean** | 0.0511 | 0.0241 | 0.0270 |
| **Standard deviation** | 0.0142 | 0.0132 | 0.0143 |
| **Hyperbolic model** | 0.0133 | 0.0078 | 0.0054 |
| **Two-tailed *p*** | <0.0001 | 0.0060 | 0.0019 |
| **Power-law model** | 0.0391 | 0.0213 | 0.0174 |
| **Two-tailed *p*** | 0.0350 | 0.5428 | 0.0804 |

Two-tailed *p*-values were calculated by comparing the empirical grand mean to each model's prediction using a *t*-test with 8 df.

The power-law model has no asymptote (unlike the hyperbolic model), but it does predict that the rate of fitness increase will decelerate over time. Indeed, deceleration was clearly evident over the full 50,000 generations analysed by Wiser et al. [4]. However, the deceleration becomes less apparent as time goes on because the fitness gains become smaller, making it difficult to tell whether the changes between two successive intervals differ from one another. Using the same fitness gains predicted by the power-law model as before, we can express the predicted deceleration as the difference in gains between 40,000 and 50,000 generations and between 50,000 and 60,000 generations, which equals 0.39%. Averaged across the nine populations, the observed deceleration was -0.29% (i.e., there was a slight acceleration), but this deviation from the power-law model prediction is not significant given the variability among populations (two-tailed *p* = 0.4175).

### (c) Heterogeneity among populations in fitness

We can use the replicate assays for each population in analyses of variance to evaluate whether the fitness differences among the populations are significant, and to estimate the among-population variance component [22]. For each generation tested, we ran a two-way ANOVA with population and block as random factors (Tables S3-S5). We excluded blocks with missing values in order to fulfil a complete-block design. At all three generations, there was highly significant variation in mean fitness among the populations (all *p* < 0.0001). We found equally strong support for that variation using the rank-based, non-parametric Friedman’s method [22].

The among-population variance component, *V*_pop_, reflects the heterogeneity in mean fitness among populations that is above and beyond the variability caused by measurement noise (including block effects). We took the square root of the variance component to generate a corresponding standard deviation, σ_pop_, that is commensurate in scale to fitness (Table 3). At all three time points, σ_pop_ was between about 4.5% and 5%, so that a typical pair of populations differs in mean fitness by several percent.

**Table 3.** Among-population variance component for fitness (*V*pop) and corresponding standard deviation (σ_pop_).

| Generations | 40,000 |  | 50,000 |  | 60,000 |  |
| --- | --- | --- | --- | --- | --- | --- |
| | $V_{\text{pop}}$ | $\sigma_{\text{pop}}$ | $V_{\text{pop}}$ | $\sigma_{\text{pop}}$ | $V_{\text{pop}}$ | $\sigma_{\text{pop}}$ |
| **All nine populations** | 0.001956 | 0.0442 | 0.002005 | 0.0448 | 0.002614 | 0.0511 |
| **Excluding hypermutators** | 0.000868 | 0.0295 | 0.000694 | 0.0263 | 0.001457 | 0.0382 |
| **Also excluding Ara+1** | 0.000342 | 0.0185 | 0.000210 | 0.0145 | 0.000182 | 0.0135 |

One source of among-population variability is that three of the nine populations in our study had evolved hypermutability, such that their point-mutation rates increased by roughly 100-fold [8,13]. Both theory and prior empirical evidence indicate that these hypermutable populations should increase in fitness somewhat faster than populations with the ancestral mutation rate, at least so long as there remain some point mutations that confer fitness benefits substantially above the increased load of deleterious mutations suffered by the hypermutators [4,12,13,23]. Indeed, we see that all three hypermutator populations—Ara+3, Ara−1, and Ara−4—have higher mean fitness than any of the other six populations at all three time points tested here (Table S2). The probability of this outcome occurring by chance alone, at any given time point, is (3/9 x 2/8 x 1/7) = 0.0119. (Owing to temporal autocorrelation, such that high and low fitness populations tend to retain their relative ranks, it is inappropriate to combine probabilities across time points.) Even if we drop the hypermutator populations, the among-population variance component for fitness remains highly significant (all *p* < 0.0001) at all three generations (Tables S3-S5). However, the corresponding standard deviation, σ_pop_, falls to between 2.5% and 4% (Table 3).

Another population, Ara+1, also stands out as unusual in two respects. First, it experienced many insertions and other non-point mutations caused by increased activity of the *IS150* mobile element [24,25]. Second, the fitness trajectory for Ara+1 was noticeably lower than that of any other population [4]. Indeed, in our data, the mean fitness of Ara+1 was at least 4% lower than any other population at all three time points tested (Table S2). If the unusual activity of *IS150* in this population is causally related to its lower fitness gains—and that connection has not been proven—then it would imply that some of the IS*150*-mediated mutations moved this population into a relatively unproductive region of genotypic space, one with fewer or smaller opportunities for continued adaptation. In any case, we can remove Ara+1 from our analysis, along with the three hypermutators, and test whether the five remaining—and seemingly normal—populations vary in their fitness levels. In fact, we still see highly significant variation (all *p* < 0.0001) at all three time points (Tables S3-S5), although the standard deviation for fitness declines further to between 1% and 2% (Table 3).

### (d) Heterogeneity among populations in rates of fitness gain and deceleration

We also performed analyses of variance to assess whether the rates of fitness increase from generations 40,000 to 60,000 were homogeneous or heterogeneous across the populations, and whether the extent of deceleration over the second half of that interval relative to the first varied among the populations. These analyses required blocks with no missing values or outliers for any of the 27 samples (i.e., 9 populations at 3 time points), because they involve calculations across the generations as well as comparisons among the populations. Twenty-nine of the 42 total blocks qualified.

There is highly significant variability among the populations in the extent of their improvement between 40,000 and 60,000 generations (Table S6). The corresponding σ_pop_ is about 1.5%. In other words, a typical pair of populations differs in their fitness gains over this period by one or two percent. These results are essentially unchanged whether we include or exclude the populations that became hypermutable (Table S6), and the variation remains significant using Friedman’s non-parametric method. One might expect a negative correlation between the extent of the fitness gain and the initial fitness at the start of this interval, which would be consistent with the tendency for diminishing-returns epistasis in the LTEE [7] and many other evolution experiments [26,27,28,29,30,31]. However, using the data from Tables 1 and S2, we find that the correlation runs in the opposite direction (*r* = 0.3921), owing largely to the fact that population Ara+1 had both the lowest fitness at 40,000 generations and the smallest gain between 40,000 and 60,000 generations. This association provides further evidence that Ara+1 has moved into a region of genotypic space with less potential for sustained adaptation. If we exclude this population, the correlation becomes negative, as expected, although it is not significant (*r* = −0.0623, 6 df, one-tailed *p* = 0.4417).

We also find significant variation among populations in the extent to which their fitness trajectories decelerated or accelerated (i.e., *W*50k/*W*40k – *W*60k/*W*50k) in the two successive 10,000-generation intervals (Table S7). This variability in the curvature of the trajectories is again significant whether or not the hypermutable populations are included, and it remains significant using Friedman’s rank-based method.

We considered the possibility that a population that was not a hypermutator at 50,000 generations might have evolved hypermutability between 50,000 and 60,000 generations. In particular, population Ara+4 had the largest fitness gain of any population between 50,000 and 60,000 generations, and it also showed the most acceleration over that period compared to the prior 10,000 generations (Table 1). We performed fluctuation tests [18] using clones from Ara+4 at 60,000 generations to assess whether their mutation rate was elevated, using the LTEE ancestral strain and clones from a more typical population, Ara+5, as comparisons. These tests show no indication of an elevated mutation rate in Ara+4 (Table S8), whereas the populations that previously evolved hypermutability showed ~100-fold increases in their mutation rates [8,13].

## 4. Discussion

Most natural populations live in environments that change often as the result of coevolving species, abiotic perturbations, or both. This environmental variability may prevent natural populations from reaching their adaptive limits. However, it is also worthwhile examining whether a population’s mean fitness can increase indefinitely, and potentially without limit, in a constant environment. Doing so may provide new insights into the limits of adaptation by natural selection. Also, while an organism’s overall fitness may be subject to changing environments, it is possible that some aspects of its performance—such as core metabolic processes—might experience constant pressures that would favour tiny improvements even after eons of selection.

The bacteria in the LTEE live and evolve in a deliberately simple and uniform environment. Except for the daily fluctuations produced by the transfers, the exogenous environment is kept as constant as feasible by using a chemically defined medium and simple protocols. The limiting resource, glucose, is provided at a low concentration by laboratory standards, resulting in cell densities and levels of secreted metabolites that are likewise low, thereby reducing, but not eliminating, opportunities for complex frequency-dependent interactions. Although some frequency-dependent interactions have evolved in the LTEE, including a transiently stable polymorphism recently discovered in one of the populations used in our study [6], there is no reason to expect that such interactions should lead to systematic increases in fitness relative to a distant ancestor or other reference competitor. That is, any context-specific interactions should favour mutations that are beneficial relative to an organism’s immediate competitors and predecessors. By contrast, fitness gains relative to a distant ancestor or other reference competitor indicate improvements in the shared, constant aspects of the environment.

Wiser et al. [4] found that a simple, two-parameter power-law model—where the rate of improvement declines over time, but fitness has no upper bound—provided an excellent description of the grand-mean fitness trajectory in the LTEE over 50,000 generations. They also showed that if the dataset was truncated, the power-law model still predicted the future trajectory with impressive accuracy. In this study, we extended the duration of that prior study by “only” 10,000 generations (i.e., 20%). However, the changes in fitness are much smaller, and the curvature in the fitness trajectory is much more subtle, over this period than during the experiment as a whole. Therefore, to ensure that our study had the power to detect the predicted changes, we obtained about 40 fitness estimates for each population at each of 40,000, 50,000 and 60,000 generations.

Our data and analyses provide strong, albeit imperfect, support for the power-law model. The mean fitness of the populations, individually and collectively, continued to improve significantly even over these later generations (Figure 1, Table 1). The average increase in relative fitness was reasonably consistent with the predictions of the power-law model based entirely on the previous data, although the difference between the 5.1% increase we observed and the predicted gain of 3.9% was marginally significant (Table 2). In any case, the fit of the power-law model was much better than that of the hyperbolic model, which predicted only a 1.3% increase in fitness (Table 2). The experimental data also showed no clear deceleration in the rate of fitness increase from 50,000 to 60,000 generations relative to that from 40,000 to 50,000 generations; however, the predicted difference between these periods is small, and the observed and predicted values do not differ significantly. In any case, if an alternative to the power-law model were to be sought based on our data, it would have even less tendency toward deceleration, not more—in other words, our new data, like the data analysed by Wiser et al. [4], do not support a model in which fitness has an upper bound.

The extensive replication in our study also enabled us to quantify the divergence of the evolving populations’ fitness trajectories. The among-population variation in mean fitness provides information on the structure of the fitness landscape that the grand-mean fitness trajectory cannot [11,32,33]. In particular, this variability sheds light on the form of epistasis, which is a key feature of the dynamic model developed in Wiser et al. [4] that gives rise to the power-law relationship. In some respects, it was already known that the fitness trajectories are not strictly parallel. First, one population evolved the ability to grow on citrate [9,10], a resource in the medium of the LTEE that remains unavailable to the other populations. This population was excluded from our analysis (and, after the function evolved, from Wiser et al. [4]) because the strong density- and frequency-dependent effects of that phenotype make the assays used to measure fitness inappropriate. (Two other populations were excluded from both studies because, in later generations, they do not make colonies on the plates used to enumerate competitors in the fitness assays.) Second, Wiser et al. [4] showed that hypermutable populations had faster-rising fitness trajectories than the other populations. However, they did not examine whether there was variation in fitness among the populations that retained the low ancestral mutation rate throughout the LTEE. Third, there was significant among-population fitness variation in the early generations of the LTEE [11,34]; however, it was not known whether it would persist or, alternatively, diminish if the populations converged on the same fitness level over time.

In this study, we found significant among-population variation in fitness at all three late-generation time points tested (Tables S3-S5). Moreover, this variation was significant even if those populations that became hypermutable were excluded from the analysis. The square root of the among-population variance component (i.e., comparable to a standard deviation) over the period from 40,000 to 60,000 generations was about 5% for all of the populations in our analysis and about 3% without the hypermutable populations (Table 3). Population Ara+1 was a notable outlier among the non-hypermutable populations, having the lowest fitness of all populations at each generation we tested; this population also had the lowest trajectory in the study by Wiser et al. [4].

Given the empirical and theoretical support for pervasive diminishing-returns epistasis in the LTEE [4,7], one would expect the Ara+1 population to show a propensity to improve faster than the other non-hypermutable populations. That is, its low fitness implies greater scope for improvement under the assumption that all populations are subject to the same form and strength of diminishing-returns epistasis. In fact, however, Ara+1 had the least improvement from 40,000 to 60,000 generations of any population—in other words, the opposite of that expectation. This result implies that this population has, in some sense, gotten stuck in a genotypic region of the fitness landscape that constrains its evolvability. It follows that the coefficient that describes the strength of diminishing-returns epistasis is not a constant—even in the constant environment of the LTEE—but instead it must vary between local neighbourhoods in genotypic space.

Besides its unique fitness trajectory, population Ara+1 also had an unusually high number of mutations caused by the IS*150* mobile element [24,25]. In some respects, that would seem to make Ara+1 similar to the populations that became hypermutable and that had the fastest rates of fitness improvement. Indeed, some genes that acquired mutations caused by IS elements in one population accumulated point mutations in other populations, implying that knockout or knockdown mutations in those genes were beneficial [35,36], which has been confirmed in some cases [37]. However, insertions are typically more disruptive of gene functions than are point mutations. Therefore, a population that evolves insertion-mediated hypermutability may have fewer opportunities for subtle refinements that could compensate for earlier mutations that, while beneficial, overshot some optimum effect. We emphasize this connection between IS-mediated hypermutability and reduced evolvability is, at present, merely conjecture. However, we think it is a hypothesis worthy of further exploration and testing.

How rugged is the fitness landscape on which the LTEE populations are evolving? De Visser and Krug [38] present several metrics that describe the ruggedness of fitness landscapes based on the individual and combined effects of mutations found in evolved genotypes. Using data from the first five mutations to fix in one population [7], the LTEE landscape appears smooth when compared to other studies in which mutations were combined and their fitness effects quantified. Smoother landscapes presumably tend to promote more repeatable adaptive trajectories and, indeed, many studies have identified parallel phenotypic and genetic changes in the LTEE [34,36,37,39,40,41]. Nonetheless, there are many examples of divergence as well. Some of these seem minor, such as the fact that very few mutations are identical at the sequence level, even when the same gene has beneficial mutations in most or all of the LTEE populations [36]. Other cases of divergence have more obvious and important consequences, such as the evolution of hypermutability in some populations [4,8]. The most striking case of divergence in the LTEE is the new ability to grow on the citrate that is present in the medium, which evolved in only one population [9,10]. In this study, we demonstrated additional divergences impacting the fitness trajectories of the seemingly ordinary populations that did not become hypermutable or evolve the ability to consume citrate. We showed that population Ara+1 is peculiar in having both the lowest and shallowest fitness trajectory. Even among the other non-hypermutable populations there is significant variation in their fitness levels. Hence, subtle divergences are ubiquitous even in the most ordinary populations. In closing, both adaptation and divergence are continuing unabated in the LTEE, even after many tens of thousands of generations in a constant environment.

Authors’ contributions. R.E.L. and M.J.W. conceived and designed the experiments. M.J.W., N.R., Z.D.B., J.R.N., J.J.M., L.Z., C.B.T., B.D.W., R.M., A.R.B., E.J.B., J.B., N.A.G., K.J.C., M.R., K.W., S.E.P., R.S., C.C., J.S.M., and N.H. performed the experiments. R.E.L. and M.J.W. analysed the data. R.E.L., M.J.W., and Z. D.B. wrote the paper; all authors approved the submitted version.

## Acknowledgements.

We thank everyone who has participated in and sustained the LTEE over the years. This research was supported, in part, by an NSF grant (DEB-1451740] and by the BEACON Center for the Study of Evolution in Action (NSF Cooperative Agreement DBI-0939454).

---

**Table S1.** The *E. coli* population samples and strains used in this study.

| **ID Number** | **Description** |
| --- | --- |
| REL10999 | Whole-population sample from Ara+1 at generation 40,000 |
| REL10933 | Whole-population sample from Ara+2 at generation 40,000 |
| REL10934 | Whole-population sample from Ara+3 at generation 40,000 |
| REL10935 | Whole-population sample from Ara+4 at generation 40,000 |
| REL10977 | Whole-population sample from Ara+5 at generation 40,000 |
| REL10926 | Whole-population sample from Ara−1 at generation 40,000 |
| REL10929 | Whole-population sample from Ara−4 at generation 40,000 |
| REL10930 | Whole-population sample from Ara−5 at generation 40,000 |
| REL10998 | Whole-population sample from Ara−6 at generation 40,000 |
| REL11383 | Whole-population sample from Ara+1 at generation 50,000 |
| REL11325 | Whole-population sample from Ara+2 at generation 50,000 |
| REL11326 | Whole-population sample from Ara+3 at generation 50,000 |
| REL11327 | Whole-population sample from Ara+4 at generation 50,000 |
| REL11362 | Whole-population sample from Ara+5 at generation 50,000 |
| REL11318 | Whole-population sample from Ara−1 at generation 50,000 |
| REL11321 | Whole-population sample from Ara−4 at generation 50,000 |
| REL11322 | Whole-population sample from Ara−5 at generation 50,000 |
| REL11382 | Whole-population sample from Ara−6 at generation 50,000 |
| REL11765 | Whole-population sample from Ara+1 at generation 60,000 |
| REL11686 | Whole-population sample from Ara+2 at generation 60,000 |
| REL11687 | Whole-population sample from Ara+3 at generation 60,000 |
| REL11688 | Whole-population sample from Ara+4 at generation 60,000 |
| REL11723 | Whole-population sample from Ara+5 at generation 60,000 |
| REL11678 | Whole-population sample from Ara−1 at generation 60,000 |
| REL11681 | Whole-population sample from Ara−4 at generation 60,000 |
| REL11682 | Whole-population sample from Ara−5 at generation 60,000 |
| REL11763 | Whole-population sample from Ara−6 at generation 60,000 |
| REL10948 | Clone A sampled from Ara−5 at generation 40,000 |
| REL11638 | Spontaneous Ara<sup>+</sup> mutant derived from REL10948 |
| REL606 | Ancestral strain for the long-term evolution experiment |
| REL11709 | Clone A sampled from Ara+4 at generation 60,000 |
| REL11710 | Clone B sampled from Ara+4 at generation 60,000 |
| REL11711 | Clone C sampled from Ara+4 at generation 60,000 |
| REL11728 | Clone A sampled from Ara+5 at generation 60,000 |
| REL11729 | Clone B sampled from Ara+5 at generation 60,000 |
| REL11730 | Clone C sampled from Ara+5 at generation 60,000 |

**Table S2.** Estimated mean fitness of nine LTEE populations at three time points, measured relative to a common competitor isolated from one population at 40,000 generations.

| Population | <u>40,000 Generations</u> |  |  | <u>50,000 Generations</u> |  |  | <u>60,000 Generations</u> |  |  |
| --- | --- | --- | --- | --- | --- | --- | --- | --- | --- |
|  | N | Mean | SD | N | Mean | SD | N | Mean | SD |
| **Ara+1** | 41 | 0.9467 | 0.0369 | 39 | 0.9682 | 0.0342 | 42 | 0.9708 | 0.0202 |
| **Ara+2** | 41 | 1.0253 | 0.0288 | 42 | 1.0388 | 0.0581 | 39 | 1.0739 | 0.0525 |
| **Ara+3** | 41 | 1.0933 | 0.0367 | 42 | 1.1251 | 0.0387 | 39 | 1.1449 | 0.0500 |
| **Ara+4** | 42 | 0.9999 | 0.0264 | 42 | 1.0079 | 0.0266 | 41 | 1.0598 | 0.0366 |
| **Ara+5** | 41 | 0.9870 | 0.0281 | 41 | 1.0164 | 0.0249 | 42 | 1.0422 | 0.0372 |
| **Ara-1** | 42 | 1.0387 | 0.0471 | 42 | 1.0846 | 0.0604 | 41 | 1.1145 | 0.0592 |
| **Ara-4** | 41 | 1.0584 | 0.0341 | 41 | 1.0661 | 0.0510 | 41 | 1.1058 | 0.0377 |
| **Ara-5** | 41 | 0.9909 | 0.0204 | 42 | 1.0256 | 0.0221 | 41 | 1.0447 | 0.0261 |
| **Ara-6** | 42 | 1.0217 | 0.0194 | 42 | 1.0383 | 0.0262 | 42 | 1.0588 | 0.0215 |
| **Grand mean** | 9 | 1.0180 | 0.0432 | 9 | 1.0412 | 0.0459 | 9 | 1.0684 | 0.0504 |

**Table S3.** ANOVAs testing among-population variation in fitness at 40,000 generations.

**All nine populations**
| **Source** | **SS** | **df** | **MS** | **F** | **p** | **Friedman p** |
| --- | --- | --- | --- | --- | --- | --- |
| **Population** | 0.58552 | 8 | 0.07319 | 89.618 | <0.0001 | <0.0001 |
| **Block** | 0.09356 | 36 | 0.00260 | 3.182 | <0.0001 |  |
| **Error** | 0.23521 | 288 | 0.00082 |  |  |  |
| **Total** | 0.91429 | 332 | 0.00275 |  |  |  |

**Excluding three hypermutator populations**
| **Source** | **SS** | **df** | **MS** | **F** | **p** | **Friedman p** |
| --- | --- | --- | --- | --- | --- | --- |
| **Population** | 0.16364 | 5 | 0.03273 | 52.091 | <0.0001 | <0.0001 |
| **Block** | 0.04599 | 36 | 0.00128 | 2.033 | <0.0001 |  |
| **Error** | 0.11310 | 180 | 0.00063 |  |  |  |
| **Total** | 0.32273 | 221 | 0.00146 |  |  |  |

**Also excluding population Ara+1**
| **Source** | **SS** | **df** | **MS** | **F** | **p** | **Friedman p** |
| --- | --- | --- | --- | --- | --- | --- |
| **Population** | 0.05289 | 4 | 0.01322 | 23.899 | <0.0001 | <0.0001 |
| **Block** | 0.03058 | 36 | 0.00085 | 1.535 | 0.0408 |  |
| **Error** | 0.07966 | 144 | 0.00055 |  |  |  |
| **Total** | 0.16313 | 184 | 0.00089 |  |  |  |

**Table S4.** ANOVAs testing among-population variation in fitness at 50,000 generations.

**All nine populations**
| **Source** | **SS** | **df** | **MS** | **F** | **p** | **Friedman p** |
| --- | --- | --- | --- | --- | --- | --- |
| **Population** | 0.60393 | 8 | 0.07549 | 58.312 | <0.0001 | <0.0001 |
| **Block** | 0.11506 | 36 | 0.00320 | 2.469 | <0.0001 |  |
| **Error** | 0.37284 | 288 | 0.00129 |  |  |  |
| **Total** | 1.09183 | 332 | 0.00329 |  |  |  |

**Excluding three hypermutator populations**
| **Source** | **SS** | **df** | **MS** | **F** | **p** | **Friedman p** |
| --- | --- | --- | --- | --- | --- | --- |
| **Population** | 0.13308 | 5 | 0.02662 | 27.859 | <0.0001 | <0.0001 |
| **Block** | 0.05954 | 36 | 0.00165 | 1.731 | 0.0106 |  |
| **Error** | 0.17196 | 180 | 0.00096 |  |  |  |
| **Total** | 0.36458 | 221 | 0.00165 |  |  |  |

**Also excluding population Ara+1**
| **Source** | **SS** | **df** | **MS** | **F** | **p** | **Friedman p** |
| --- | --- | --- | --- | --- | --- | --- |
| **Population** | 0.03505 | 4 | 0.00876 | 8.796 | <0.0001 | <0.0001 |
| **Block** | 0.04746 | 36 | 0.00132 | 1.323 | 0.1269 |  |
| **Error** | 0.14346 | 144 | 0.00100 |  |  |  |
| **Total** | 0.22597 | 184 | 0.00123 |  |  |  |

**Table S5.**
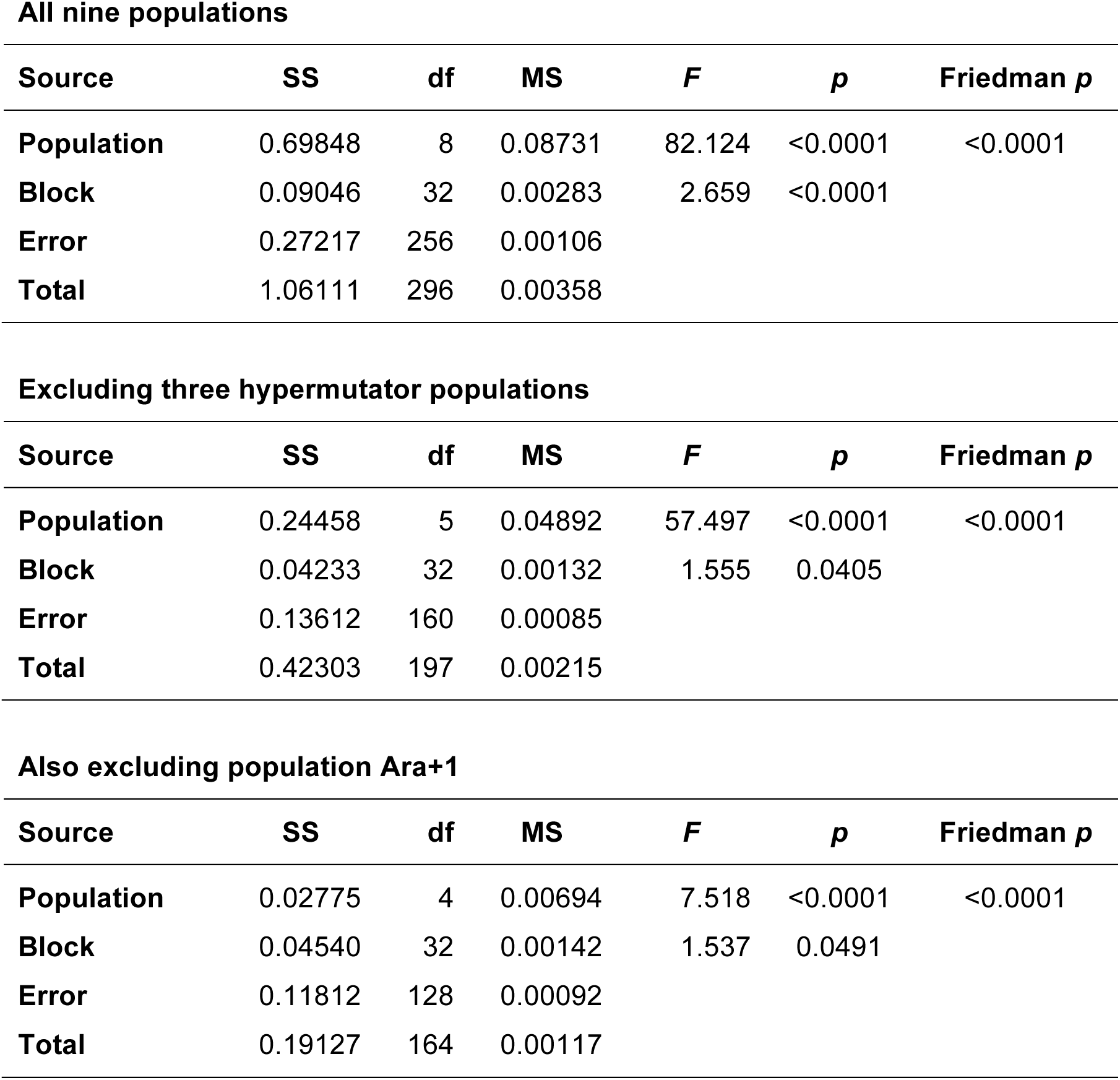
ANOVAs testing among-population variation in fitness at 60,000 generations.

**All nine populations**
| Source | SS | df | MS | *F* | *p* | Friedman *p* |
| --- | --- | --- | --- | --- | --- | --- |
| Population | 0.69848 | 8 | 0.08731 | 82.124 | <0.0001 | <0.0001 |
| Block | 0.09046 | 32 | 0.00283 | 2.659 | <0.0001 |  |
| Error | 0.27217 | 256 | 0.00106 |  |  |  |
| Total | 1.06111 | 296 | 0.00358 |  |  |  |

**Excluding three hypermutator populations**
| Source | SS | df | MS | *F* | *p* | Friedman *p* |
| --- | --- | --- | --- | --- | --- | --- |
| Population | 0.24458 | 5 | 0.04892 | 57.497 | <0.0001 | <0.0001 |
| Block | 0.04233 | 32 | 0.00132 | 1.555 | 0.0405 |  |
| Error | 0.13612 | 160 | 0.00085 |  |  |  |
| Total | 0.42303 | 197 | 0.00215 |  |  |  |

**Also excluding population Ara+1**
| Source | SS | df | MS | *F* | *p* | Friedman *p* |
| --- | --- | --- | --- | --- | --- | --- |
| Population | 0.02775 | 4 | 0.00694 | 7.518 | <0.0001 | <0.0001 |
| Block | 0.04540 | 32 | 0.00142 | 1.537 | 0.0491 |  |
| Error | 0.11812 | 128 | 0.00092 |  |  |  |
| Total | 0.19127 | 164 | 0.00117 |  |  |  |

**Table S6.** ANOVAs testing among-population variation in fitness gains between 40,000 and 60,000 generations.

**All nine populations**
| **Source** | **SS** | **df** | **MS** | ***F*** | ***p*** | **Friedman *p*** |
| --- | --- | --- | --- | --- | --- | --- |
| **Population** | 0.05047 | 8 | 0.00631 | 4.353 | 0.0001 | <0.0001 |
| **Block** | 0.05595 | 28 | 0.00200 | 1.379 | 0.1055 |  |
| **Error** | 0.32460 | 224 | 0.00145 |  |  |  |
| **Total** | 0.43102 | 260 | 0.00166 |  |  |  |

**Excluding three hypermutator populations**
| **Source** | **SS** | **df** | **MS** | ***F*** | ***p*** | **Friedman *p*** |
| --- | --- | --- | --- | --- | --- | --- |
| **Population** | 0.04233 | 5 | 0.00847 | 7.277 | <0.0001 | <0.0001 |
| **Block** | 0.02563 | 28 | 0.00092 | 0.787 | 0.7667 |  |
| **Error** | 0.16286 | 140 | 0.00116 |  |  |  |
| **Total** | 0.23082 | 173 | 0.00133 |  |  |  |

**Table S7.**
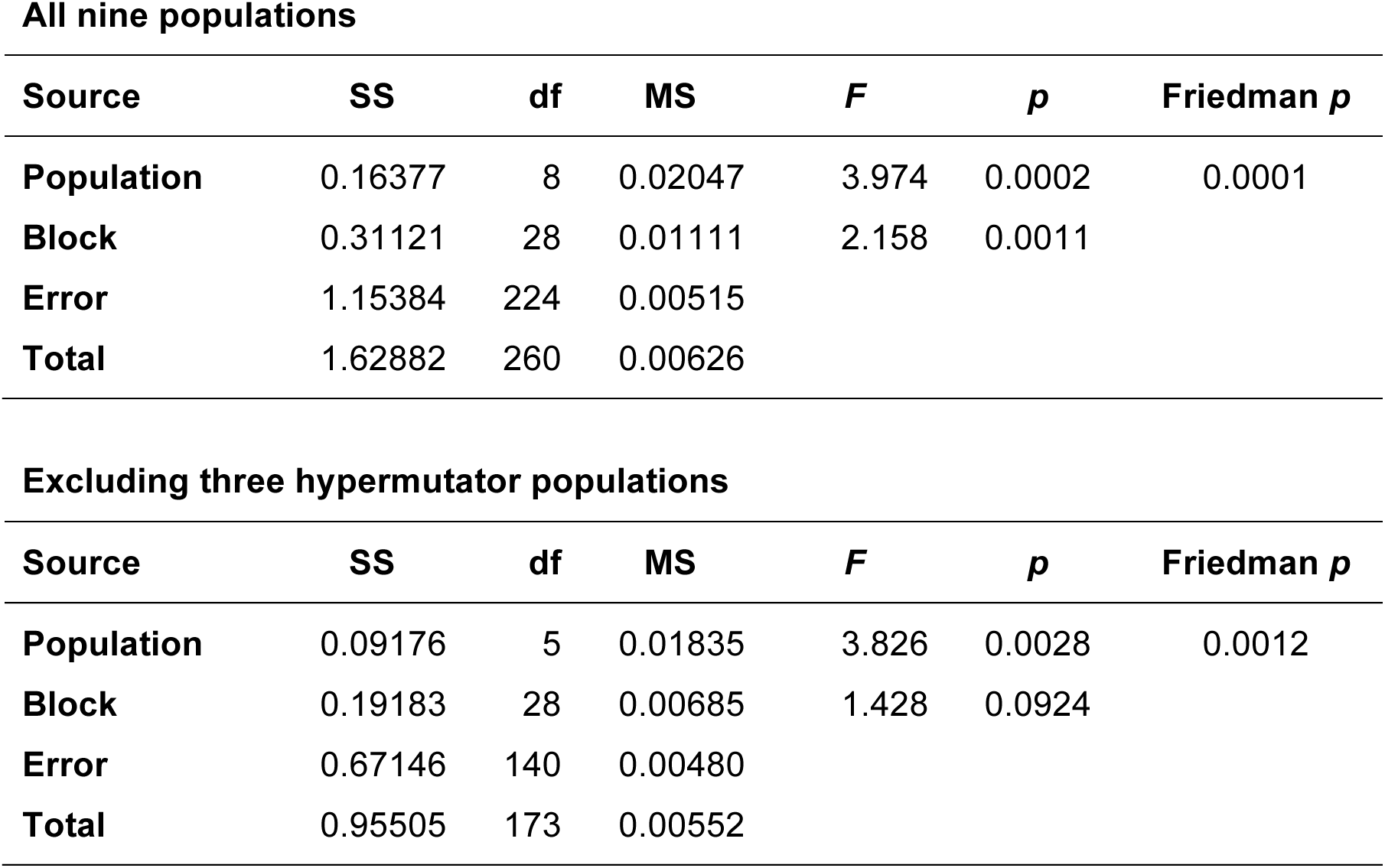
ANOVAs testing among-population variation in the deceleration of fitness changes between 40,000 to 50,000 and 50,000 to 60,000 generations.

**All nine populations**
| Source | SS | df | MS | *F* | *p* | Friedman *p* |
| --- | --- | --- | --- | --- | --- | --- |
| Population | 0.16377 | 8 | 0.02047 | 3.974 | 0.0002 | 0.0001 |
| Block | 0.31121 | 28 | 0.01111 | 2.158 | 0.0011 |  |
| Error | 1.15384 | 224 | 0.00515 |  |  |  |
| Total | 1.62882 | 260 | 0.00626 |  |  |  |

**Excluding three hypermutator populations**
| Source | SS | df | MS | *F* | *p* | Friedman *p* |
| --- | --- | --- | --- | --- | --- | --- |
| Population | 0.09176 | 5 | 0.01835 | 3.826 | 0.0028 | 0.0012 |
| Block | 0.19183 | 28 | 0.00685 | 1.428 | 0.0924 |  |
| Error | 0.67146 | 140 | 0.00480 |  |  |  |
| Total | 0.95505 | 173 | 0.00552 |  |  |  |

**Table S8.** Fluctuation tests to estimate mutation rates using the *p*_0_ method.

| Strain | Population /<br>Generation | Average<br>Cells Tested | Replicates | Mutation<br>Events | Estimated<br>Mutation Rate |
| --- | --- | --- | --- | --- | --- |
| REL606 | Ancestor / 0 | $9.41 \times 10^7$ | 24 | 6 | $3.06 \times 10^{-9}$ |
| REL11709 | Ara+4 / 60,000 | $2.92 \times 10^7$ | 24 | 1 | $1.46 \times 10^{-9}$ |
| REL11710 | Ara+4 / 60,000 | $2.32 \times 10^7$ | 24 | 1 | $1.84 \times 10^{-9}$ |
| REL11711 | Ara+4 / 60,000 | $2.56 \times 10^7$ | 24 | 0.5 | $8.24 \times 10^{-10}$ |
| REL11728 | Ara+5 / 60,000 | $3.16 \times 10^7$ | 24 | 1 | $1.35 \times 10^{-9}$ |
| REL11729 | Ara+5 / 60,000 | $5.37 \times 10^7$ | 24 | 2 | $1.62 \times 10^{-9}$ |
| REL11730 | Ara+5 / 60,000 | $5.10 \times 10^7$ | 24 | 2 | $1.71 \times 10^{-9}$ |

**Mutations to nalidixic-acid resistance**
| Strain | Population /<br>Generation | Average<br>Cells Tested | Replicates | Mutation<br>Events | Estimated<br>Mutation Rate |
| --- | --- | --- | --- | --- | --- |
| REL606 | Ancestor / 0 | $9.41 \times 10^7$ | 24 | 1 | $4.52 \times 10^{-10}$ |
| REL11709 | Ara+4 / 60,000 | $2.92 \times 10^7$ | 24 | 3 | $4.57 \times 10^{-9}$ |
| REL11710 | Ara+4 / 60,000 | $2.32 \times 10^7$ | 24 | 1 | $1.84 \times 10^{-9}$ |
| REL11711 | Ara+4 / 60,000 | $2.56 \times 10^7$ | 24 | 0.5 | $8.24 \times 10^{-10}$ |
| REL11728 | Ara+5 / 60,000 | $3.16 \times 10^7$ | 24 | 0.5 | $6.66 \times 10^{-10}$ |
| REL11729 | Ara+5 / 60,000 | $5.37 \times 10^7$ | 24 | 0.5 | $3.92 \times 10^{-10}$ |
| REL11730 | Ara+5 / 60,000 | $5.10 \times 10^7$ | 24 | 0.5 | $4.13 \times 10^{-10}$ |

